# A Natural-History Study of Reproductive Scaling and Clonality in Co-Occurring *Catopsis* Griesb

**DOI:** 10.64898/2026.08.12.742214

**Authors:** Joshua M. Felton, Kaelin T. Escalante, Kamille D. Mendez, Denver T. Cayetano, Chelsea D. Specht

**Author notes:** Author for correspondence: Joshua M. Felton.

## Abstract

Multi-species communities of epiphytic bromeliads are a defining component of Neotropical canopies, yet we have little knowledge of how closely related species differ in the balance struck among vegetative growth, clonal propagation, and sexual reproduction. We compared reproductive output, clonality, and sexual systems in sympatric populations of *Catopsis nutans* (Sw.) Griseb. and *Catopsis sessiliflora* (Ruiz & Pav.) Mez occupying citrus groves in central Belize, sampling 235 reproductively mature individuals across 138 host trees in nine groves. Reproductive output, measured as the count of reproductive structures per individual, was modeled with negative binomial generalized linear mixed models that accounted for vegetative size and host tree identity. All sampled *C. nutans* were hermaphroditic, whereas all *C. sessiliflora* were unisexual, representing a dioecious population. Reproductive output increased with vegetative size in both species, and the scaling relationship did not differ between them despite their differing sexual systems. After accounting for size, *C. sessiliflora* produced more reproductive structures and more connected pups than *C. nutans*, and we found no evidence of a trade-off between clonal and sexual output in both species. Within *C. sessiliflora*, staminate individuals produced more flowers than pistillate individuals across comparable sizes. Co-occurring *Catopsis* can differ markedly in baseline reproductive and clonal output while sharing a conserved scaling of output on vegetative size, offering a foundation for further comparisons of sex-specific reproductive allometry in bromeliads.

## Introduction

Bromeliaceae are a globally significant component of Neotropical forest biodiversity, contributing greatly to nutrient cycling (Ladino et al., 2019), and the provisioning of habitat and resources for arthropods, birds, and other canopy fauna (Taylor et al., 2022). Epiphytic bromeliads more broadly are among the plant groups most vulnerable to tropical deforestation and the conversion of forests to agriculture and other land uses due to their reliance on intact canopy structure, stable microclimates, and long-lived host trees that take decades to develop suitable bark characteristics and water-holding capacity. Although epiphyte communities often decline sharply when forests are converted, some woody agricultural systems retain enough canopy complexity to support persistent epiphytic communities; this suggests that the effects of land-use change on epiphytes can vary depending on the type and structure of the resulting habitat (Krömer et al., 2025). Nevertheless, the reproductive ecology of epiphytic taxa in forested agricultural environments remain poorly understood. In these environments with more open and young canopies, epiphytic bromeliads may experience strong trade-offs between vegetative growth, clonal propagation, and sexual reproduction, as resources allocated toward flowering and fruit production may reduce opportunities for investment in vegetative persistence and pup production important for establishing new colonies with critical mass (Bodine et al., 2023).

*Catopsis* Griseb is an epiphytic bromeliad genus found throughout Florida, Mexico, Central America, the Antilles and South America, where many species are abundant in open canopies as twig epiphytes relying on small rosette structures that form ‘tanks’ for water impoundment (Figure1). Despite this ecological preference for sunny and exposed canopies, comparatively little is known about how closely related *Catopsis* species differ in their reproductive output, clonality, and sexual systems (Felton et al., 2026) and how these features manifest in their growth in forested agricultural fields (i.e. ‘groves’) with relatively ample space and nutrients. *Catopsis* species are particularly suitable for such comparisons because many co-occurring species exhibit substantial variation in reproductive biology, including frequent dioecy within the genus, occurring in approximately half of the described species (Martínez Correa, 2019). In dioecious species, reproductive investment frequently differs between staminate and pistillate individuals, with female function typically requiring greater resource allocation during fruit and seed production (Lloyd & Webb, 1977). These differences in physiological investment may generate sex-specific scaling relationships between vegetative size and reproductive output such that we predicted that hermaphrodites and pistillate individuals would differ in size–reproduction scaling because seed production is expected to impose greater reproductive costs.

**Figure 1.**
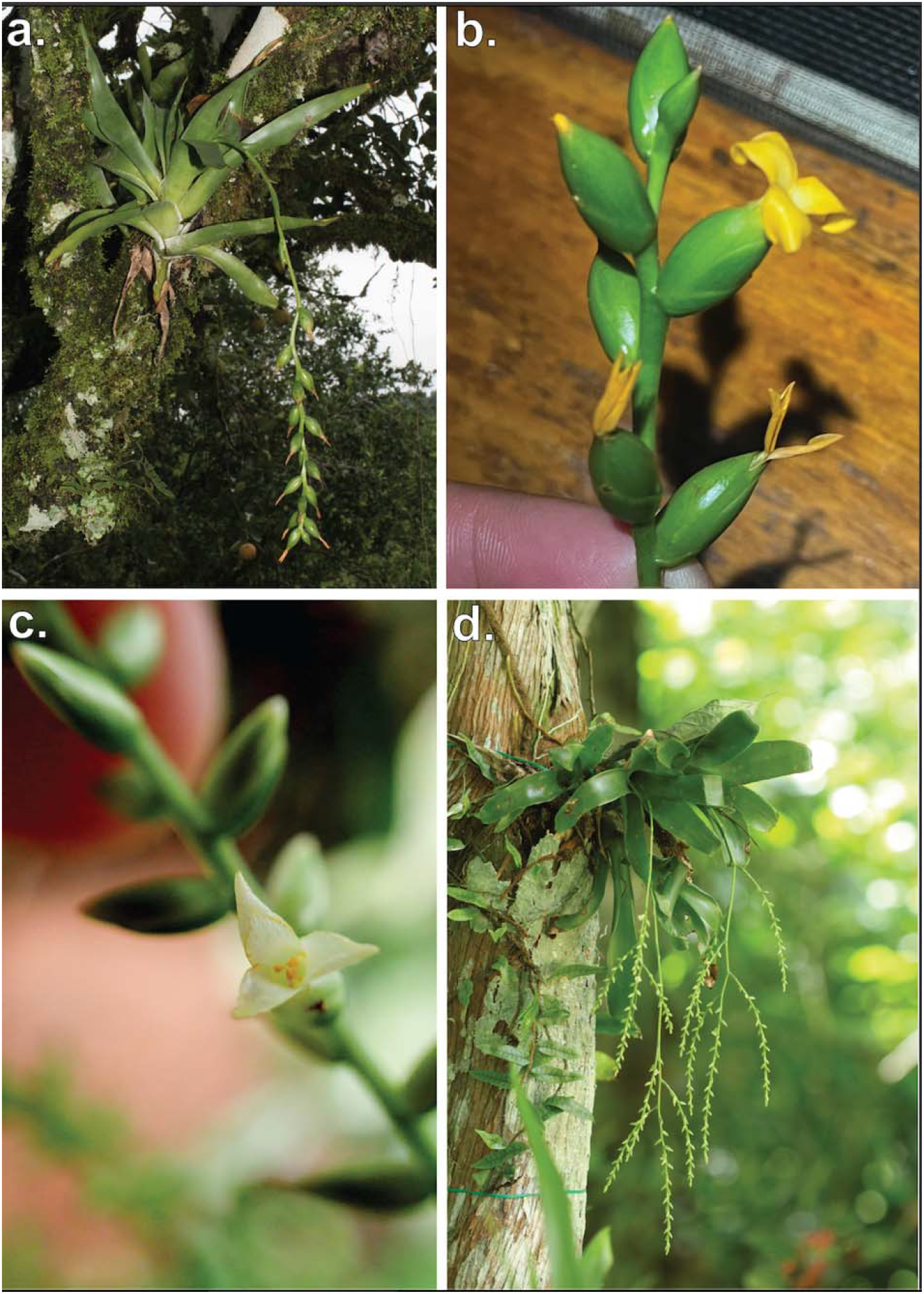
*Catopsis* growth habit and floral morphology. a–b *Catopsis nutans*: a mature rosette with a pendant, elongate inflorescence bearing showy yellow flowers; b close-up of the inflorescence showing yellow, tubular flowers subtended by green bracts. c–d *Catopsis sessiliflora*: c close-up of a solitary white flower; d rosette habit with a long, pendant, branched inflorescence bearing numerous small flowers.

Citrus groves in central Belize support sympatric populations of multiple *Catopsis* species within a structurally uniform canopy environment, making them an unusually tractable system for comparing reproductive output and vegetative architecture among closely related epiphytic taxa within a forested agricultural setting. Although citrus groves vary in epiphyte community composition depending on grove age and management practices (Krömer et al., 2025), older unmanaged groves can support epiphyte assemblages approaching forest-level richness. Here, we compared reproductive output, clonality, and sexual systems in sympatric populations of *Catopsis nutans* and *Catopsis sessiliflora* occurring within Belize citrus groves. Specifically, we asked whether (1) species differed in reproductive and clonal allocation after accounting for vegetative size, predicting that hermaphrodites would have the greatest trade-off between clonal persistence and sexual reproduction given their floral morphology (2) staminate and pistillate individuals of *C. sessiliflora* differ in reproductive output, predicting that pistillate individuals would show reduced vegetative growth or altered size-reproduction scaling relative to staminate individuals as a consequence of the greater resource demands of fruit and seed production; and (3) closely related species occupying the same agroecosystems exhibit distinct host occupancy patterns, predicting that sympatric congeners sharing a limited pool of suitable host trees would partition host use and location of persistence as a mechanism of coexistence.

## Methods

### Sampling

Fieldwork was conducted across nine citrus groves in central Belize during January 2026. Sampling included 235 reproductively mature individuals of *C. nutans* and *C. sessiliflora* distributed across 138 host trees. Individual selection was opportunistic, targeting all accessible reproductive individuals within each grove that could be safely reached from ground level or by climbing without gear. Non-reproductive individuals, and individuals that could not be safely accessed were excluded. Groves varied in size and host composition; most groves contained primarily grapefruit trees, with orange trees present in a subset of groves. Multiple individuals per host tree were sampled where present, and individuals were surveyed across multiple trees and canopy positions within each grove to characterize variation in vegetative size, reproductive state, and microhabitat occupancy. Whether the host tree was a grapefruit or orange was recorded for all sampled individuals to document patterns of host occupancy within sampled groves. Voucher specimens for all sampled species were deposited at the Belize National Herbarium (BRH; Table S1).

### Reproductive traits

Reproductive traits were measured in situ for each individual. For unisexual individuals, sex was determined based on reproductive morphology observed at the time of sampling. Staminate individuals produced pollen-bearing flowers with an immature and aborted gynoecium that lacked evidence of fruit development, whereas pistillate individuals bore developing or mature fruits. Inflorescence length was measured from the center of the vegetative axis to the distal end of the inflorescence with a flexible rod to capture true length with the curvature of most inflorescences. In individuals with paniculate inflorescences, measurements were taken to the end of the primary axis.

Reproductive output was quantified as the total count of flowers per individual. We use “reproductive output” (number of floral units produced) rather than reproductive allocation throughout as pollen v. ovary/fruit production differ substantially in per-unit resource investment over time and are not equivalent currencies for allocation (Obeso, 2002). Counts are therefore best interpreted as a proxy of reproductive output rather than direct measures of cost to the individual. For pistillate individuals, all fruits including aborted or undeveloped fruits within an infructescence were counted as evidence of having produced a flower; the terminal apical fruit, which was consistently aborted across sampled individuals, was excluded. For staminate individuals, the total number of flowers from the inflorescence that had recently flowered were counted.

### Vegetative traits

Vegetative traits were quantified to evaluate relationships among body size, clonality, and reproductive output. Longest leaf length (LLL) was used as a proxy for vegetative size and measured from the vegetative axis to the apex of the longest leaf. Clonal propagation was quantified by counting the number of pups physically connected to each focal individual.

### Microhabitat characteristics

For each epiphytic individual, microhabitat variables were recorded. Branch diameter at the point of attachment was measured to assess potential relationships between substrate size and epiphyte vegetative size. Canopy position was categorized as low (first two primary branches), middle (intermediate branches), or high (uppermost two branches) based on relative vertical position within the host canopy. These categories reflect relative canopy strata rather than absolute height. Attachment height was measured as the vertical distance from the point of attachment to ground level.

### Statistical analyses

Statistical analyses were conducted in R version 4.5.1 (R Core Team). Data processing and visualization used tidyverse v.2.0.0 (Wickham et al., 2019) and ggplot2 v.4.0.0 (Wickham, 2016). Reproductive output (flower count) was modeled using generalized linear mixed models fit with glmmTMB v.1.1.14 (Brooks et al., 2017). We first fit a Poisson model and tested for overdispersion using DHARMa v.0.5.0 (Hartig, 2026), which was significant (dispersion ratio = 1.99, P < 0.001); a negative binomial error structure was used for all subsequent models. Across species, we tested whether reproductive output varies with plant size, whether output differs between species, and whether species differ in the strength of the size–reproduction relationship (species × size interaction). Host tree identity was included as a random intercept in all models, nested within grove identity, to account for non-independence among individuals on the same host tree and among the nine sampled groves. Variance inflation factors were calculated using performance v.0.17.1 (Lüdecke et al., 2021) and indicated no problematic collinearity among fixed effects (VIF = 1.00 for both terms). Because residual diagnostics indicated substantially greater unexplained variance in *C. sessiliflora* than *C. nutans* under a shared dispersion parameter, the dispersion parameter was modeled as a function of species, which improved model fit (ΔAIC = 102.8) and resolved the residual dispersion diagnostic (DHARMa dispersion test: dispersion = 1.03, P = 0.736; zero-inflation test: P = 0.584).

To compare reproductive output between species using a comparable reproductive currency, we additionally restricted this comparison to pistillate *C. sessiliflora* individuals, since fruit and seed production in pistillate individuals is more directly comparable to the hermaphroditic reproductive structures counted in *C. nutans* than pooled staminate and pistillate output.

Within *Catopsis sessiliflora*, we tested whether reproductive output differs between staminate and pistillate individuals, whether it increases with plant size, and whether these size–reproduction relationships differ by sex, using negative binomial generalized linear mixed models. Host tree identity was included as a random intercept to account for shared host environments among co-occurring individuals.

Differences in clonal propagation (pup number) between species were evaluated using a Conway-Maxwell-Poisson generalized linear model, selected because this trait showed significant underdispersion relative to Poisson expectations (dispersion ratio = 0.53, P < 0.001). Models incorporating host tree or grove random intercepts failed to converge, consistent with limited within-group variation in this trait (90% of individuals produced 1–2 pups), so a fixed-effects model was used. The relationship between vegetative size and branch diameter was evaluated using linear models, testing whether plant size varies as a function of branch diameter and whether this relationship differs between species.

## Results

All sampled individuals of *C. nutans* (n = 124) were hermaphroditic, whereas all individuals of *C. sessiliflora* were functionally dioecious with 51 staminate and 60 pistillate individuals sampled (Table 1). Reproductive output count increased significantly with vegetative size across both focal species (negative binomial generalized linear mixed model: β = 0.092 ± 0.011, P < 0.001; Figure 2). After accounting for vegetative size, *C. sessiliflora* produced more flowers than *C. nutans* (β = 2.675 ± 0.130, P < 0.001); this difference remained significant, though attenuated, when the comparison was restricted to pistillate individuals only (β = 1.380 ± 0.112, P < 0.001). Although reproductive output increased with vegetative size in both species (*C. nutans*: β = 0.093 ± 0.012; *C. sessiliflora*: β = 0.088 ± 0.027), the interaction between species and vegetative size was not significant (likelihood ratio test: χ^2^ = 0.026, df = 1, P = 0.872), suggesting similar scaling relationships across focal taxa. In the dioecious *Catopsis sessiliflora*, staminate individuals consistently produced more flowers than pistillate individuals across comparable vegetative sizes (Figure 3; β = 2.034 ± 0.093 SE, P < 0.001). The interaction between sex and vegetative size was not significant (likelihood ratio test: χ^2^ = 0.067, df = 1, P = 0.796), indicating similar size-reproduction scaling between the unisexual flowers.

**Table 1:**
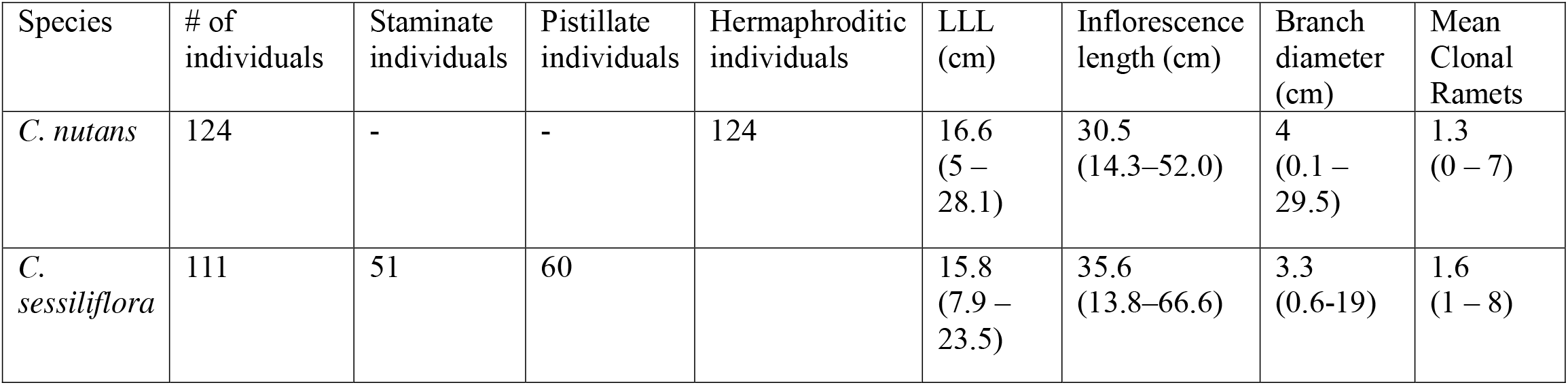
Sampling overview and vegetative characteristics of focal *Catopsis* species occurring in Belize citrus agroecosystems.

**Figure 2.**
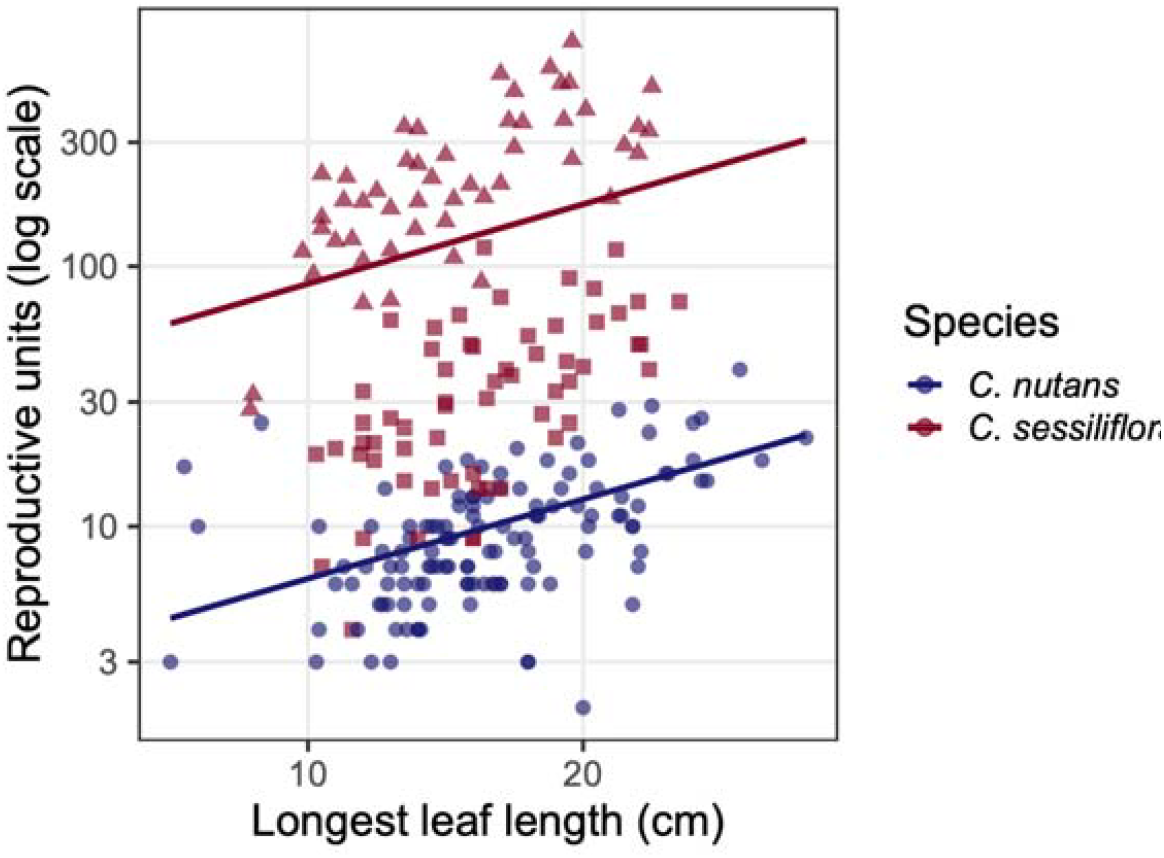
Relationship between vegetative size (longest leaf length, cm) and flowers in sympatric populations of *Catopsis nutans* (blue circles) and *C. sessiliflora* (burgundy; staminate individuals = triangles, pistillate individuals = squares) occurring within Belize citrus groves. Lines show model-predicted values from a negative binomial GLMM accounting for host tree identity. Reproductive structure counts are displayed on a log scale. Note that flowers (staminate individuals) and fruits (pistillate and hermaphroditic individuals) represent different stages of the reproductive pathway and are not equivalent currencies for allocation; counts should be interpreted as indices of reproductive output rather than direct measures of allocation cost.

**Figure 3.**
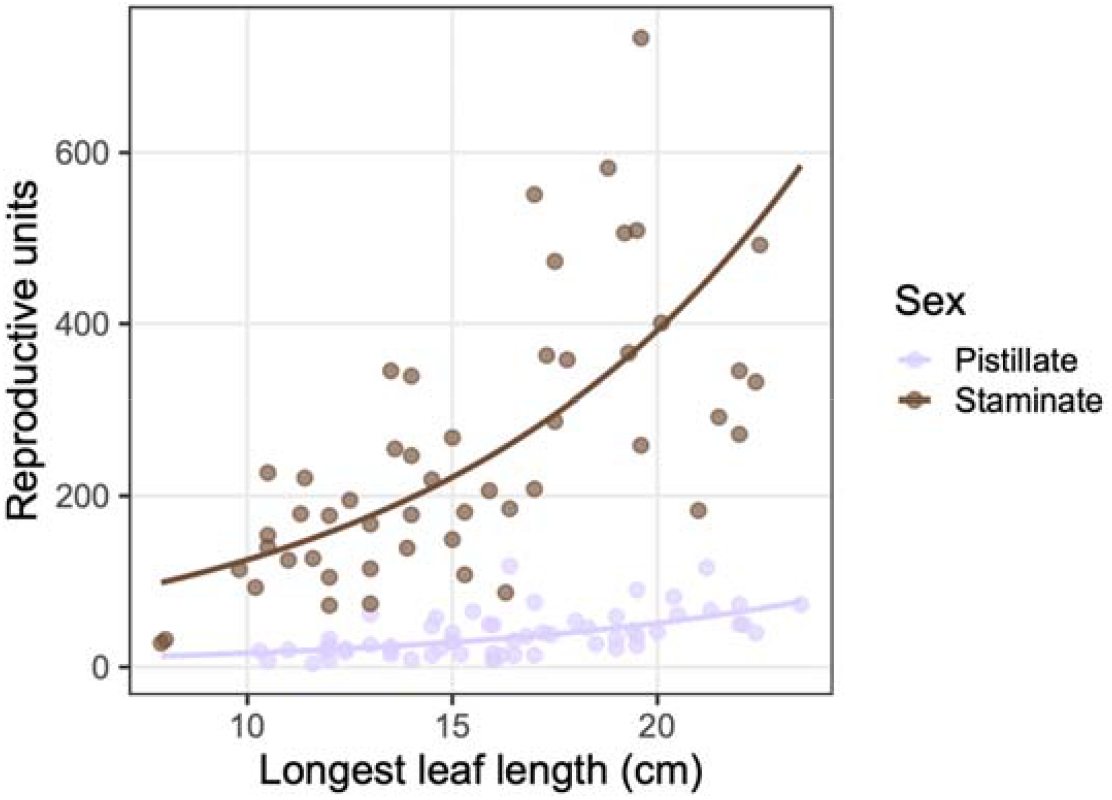
Sex-specific scaling of reproductive output with vegetative size in *Catopsis sessiliflora*. Points show individual staminate (brown) and pistillate (lavender) individuals; lines show model-predicted values from negative binomial GLMMs accounting for host tree identity. Both sexes show significant positive relationships between longest leaf length and flowers produced, but staminate individuals produce substantially more flowers across the range of observed vegetative sizes.

Clonal propagation differed between focal species (Figure 4). *Catopsis sessiliflora* produced more connected pups per individual than *C. nutans* (β = 0.246 ± 0.069, P < 0.001), indicating greater vegetative proliferation within shared citrus canopy environments. Although *C. sessiliflora* produced more connected pups than *C. nutans*, pup number showed no significant relationship with reproductive output after accounting for vegetative size, species identity, host tree, and grove (β = 0.079 ± 0.047, P = 0.093; Figure S1), and the direction of this trend was positive rather than negative, offering no support for a clonal/sexual reproduction trade-off, suggesting that clonal investment may reflect future reproductive potential rather than current flowering effort. Vegetative size was not significantly associated with host branch diameter across focal species (linear model: P = 0.63), suggesting that plant size in *C. nutans* and *C. sessiliflora* are not strongly constrained by branch-level habitat specialization (such as being a twig epiphyte) or by substrate stability (preferring larger twigs that are more stable) more broadly.

**Figure 4.**
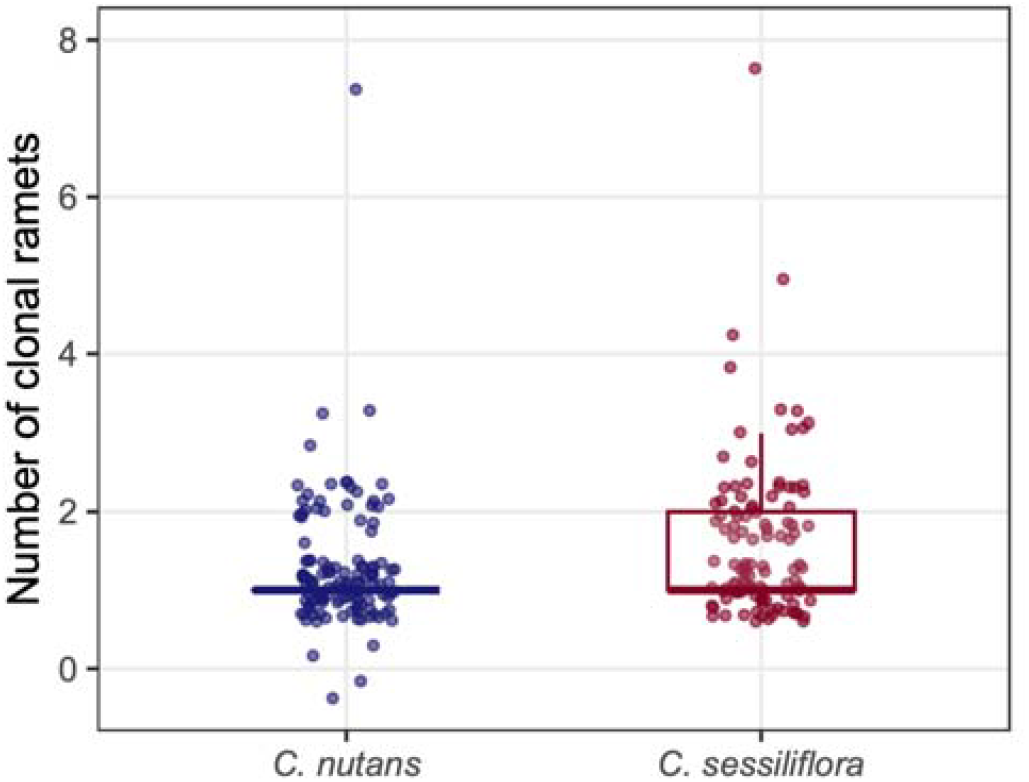
Clonal ramet production in sympatric *Catopsis nutans* and *C. sessiliflora* occurring within Belize citrus groves. Points show jittered individual observations; boxes show interquartile range with median. *Catopsis sessiliflora* produced significantly more connected pups per individual than *C. nutans* (Conway-Maxwell-Poisson generalized linear model: β = 0.246 ± 0.069, P < 0.001), indicating greater investment in vegetative proliferation despite occupying the same canopy environment

## Discussion

Woody agroecosystems are increasingly recognized as important refugia for epiphytic diversity, particularly when canopy structure and tree age allow epiphyte colonization to proceed over time (Richards et al., 2020), yet comparatively little is known about how epiphytes allocate resources toward growth, clonality, and sexual reproduction within human-modified canopy systems. Not all agricultural land uses are equally suitable. Timber monocultures with short rotation times tend to function as propagule sinks rather than sources, while structurally complex agroforestry systems with long-lived shade trees can support substantially richer epiphyte assemblages (Einzmann & Zotz, 2016). Our results demonstrate that *Catopsis* species maintain substantial reproductive and vegetative investment within Belize citrus groves while exhibiting distinct allocation patterns among species and sexes.

Across focal taxa, larger individuals consistently produced more flowers, indicating positive scaling between vegetative size and reproductive output. This pattern is often expected in modular plants, where vegetative size reflects both resource acquisition capacity and developmental timepoint, and is consistent with reproductive allometry documented across iteroparous perennial plants (Dorken et al., 2025).

Notably, the scaling relationship did not differ significantly between species despite their contrasting sexual systems, suggesting that the allometric relationship between body size and reproductive investment may be conserved in *Catopsis* regardless of whether reproduction proceeds through hermaphroditic flowers or separate staminate and pistillate individuals. Whether this reflects shared developmental constraints within *Catopsis* or a more general feature of epiphytic bromeliads remains an open question, as comparable data for other bromeliad genera are limited.

Although scaling relationships were broadly similar, *C. sessiliflora* consistently produced more flowers and more connected pups than *C. nutans* across comparable vegetative sizes, consistent with differences in resource acquisition capacity or allocation strategy. Bromeliad species differ substantially in their allocation to sexual reproduction versus clonal propagation, as well as in the timing of axillary ramet production relative to flowering (Jabaily et al., 2021). The higher reproductive and clonal output in *C. sessiliflora* may reflect either interspecific differences in life-history allocation strategies or variation in resource acquisition capacity relative to *C. nutans*. Future work should test whether these differences are associated with physiological variation in resource status through measures of foliar nitrogen content, photosynthetic rate, or other traits related to carbon and nutrient acquisition. Within *C. sessiliflora*, staminate individuals produced substantially more reproductive units than pistillate individuals across comparable vegetative sizes. The higher counts in staminate individuals may partly reflect the greater number of flowers required to achieve a given level of pollen output, rather than fundamentally greater reproductive effort. This pattern is consistent with sex allocation theory, which predicts that male fitness gain curves decelerate less steeply than female gain curves, favoring investment in a greater number of flowers to maximize pollen export (Charnov, 1979), whereas female function carries substantially higher per-unit resource costs from provisioning fruits and seeds, favoring investment in fewer, larger reproductive structures instead.

Despite theoretical expectations that investment in clonal propagation may reduce allocation toward sexual reproduction (Jabaily et al., 2021; Benzing, 2000), we detected no evidence that pup production negatively predicted reproductive output after accounting for species identity and vegetative size. This is consistent with the acquisition-allocation framework of Van Noordwijk & De Jong (1986) which shows that individual variation in total resource acquisition can produce positive correlations between traits even when a genuine allocation trade-off exists at the level of a fixed resource pool. Jabaily et al. (2021) similarly found that axillary ramet production did not consistently reduce parental ramet growth rates in bromeliads, suggesting that variation in total resource acquisition or vegetative vigor among individuals may drive allocation trade-offs, rather than these trade-offs reflecting fixed, species-level life-history strategies.

Several natural-history observations from Belize citrus groves suggest ecological differences among the focal taxa despite broad overlap in canopy occupation. Although our surveys were not designed to test host preference, *C. nutans* was observed only on orange trees and not on grapefruit trees in groves where both host species were present. Because we did not quantify host availability or sampling effort by tree species, this pattern should be regarded as an anecdotal natural-history observation rather than evidence of host preference. Both species occupied the same canopy strata (low and high), although *C. sessiliflora* occurred proportionally more often in the upper canopy than *C. nutans* (Fisher’s exact test: P = 0.017; Figure S3). This overlap in occupied habitat, despite the significant difference in relative frequency, likely contributes to field identification difficulty alongside morphological similarity. In particular, infructescences of *C. sessiliflora* consistently became pendant during fruiting and frequently remained pendant following anthesis in staminate individuals, suggesting that scape orientation may vary substantially across developmental stages and should be used cautiously as a diagnostic character. Similarity in vegetative morphology and shared occupancy of exposed twig habitats may further contribute to misidentifications within the genus, particularly in community science observations and herbarium collections.

Taken together, these patterns highlight substantial variation in reproductive strategy among co-occurring *Catopsis* species. Further work examining carbon and nitrogen content of reproductive versus vegetative tissues would allow more direct comparison with the resource-based frameworks that have proven productive in studies of other dioecious plants (Harris & Pannell, 2008) and would help establish whether the patterns observed here reflect general principles of epiphyte reproductive ecology or peculiarities of *Catopsis*.

## Supporting information

Table S1 BRH vouchers

## Author Contributions

JMF: Conceptualization, Methodology, Investigation, Writing – original draft, Visualization KTE: Investigation, Writing – original draft KDM: Investigation, Writing – original draft DTC: Writing – final draft CDS: Writing – review & editing

## Acknowledgments

We thank the University of Belize Environmental Research Institute for their generosity in supplying field equipment, and are grateful to Ella Baron, John Gregorio, Edwin Miranda, Wilfredo Miranda, Jorge Aldana and Pascual Garcia of lan Anderson’s Lodge Caves Branch Botanical Garden for their assistance in the field and funding.

Photo credits: a, David Amaya; b, Joshua Felton; c, Ella Barron; d, Bruce Holst.

---

**Figure S1.**
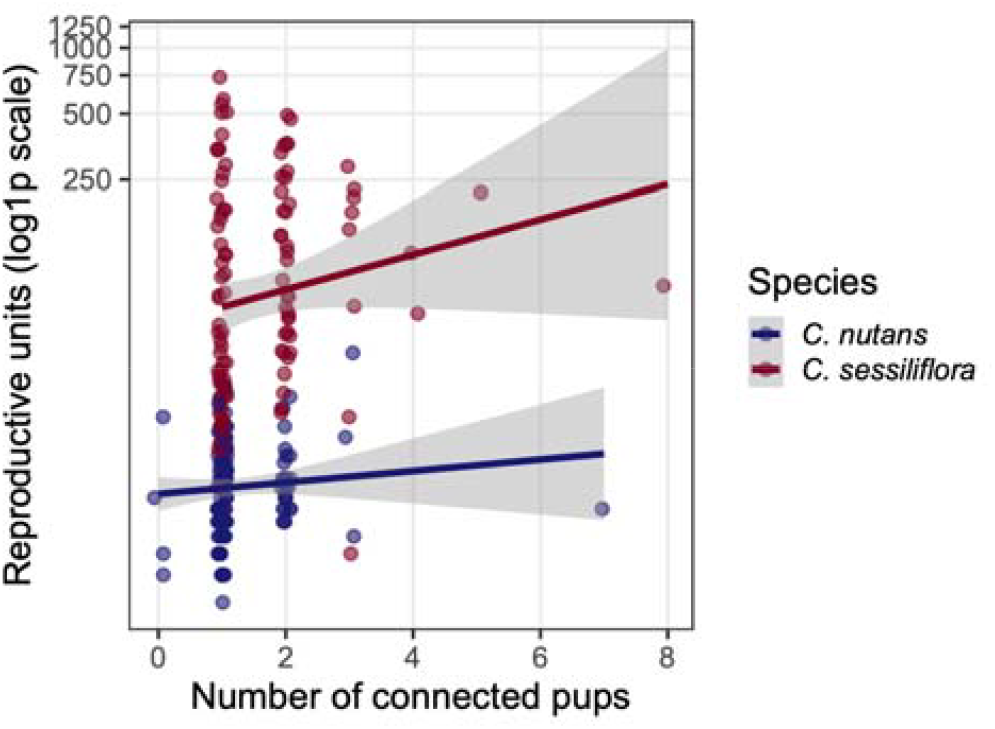
Relationship between clonal propagation and reproductive output in focal *Catopsis* species. Reproductive output, measured as the number of reproductive units per individual, was not significantly associated with the number of connected pups after accounting for vegetative size and species identity. Points represent individual plants and lines show linear trends for visualization. Y-axis shown on a log1p

**Figure S2.**
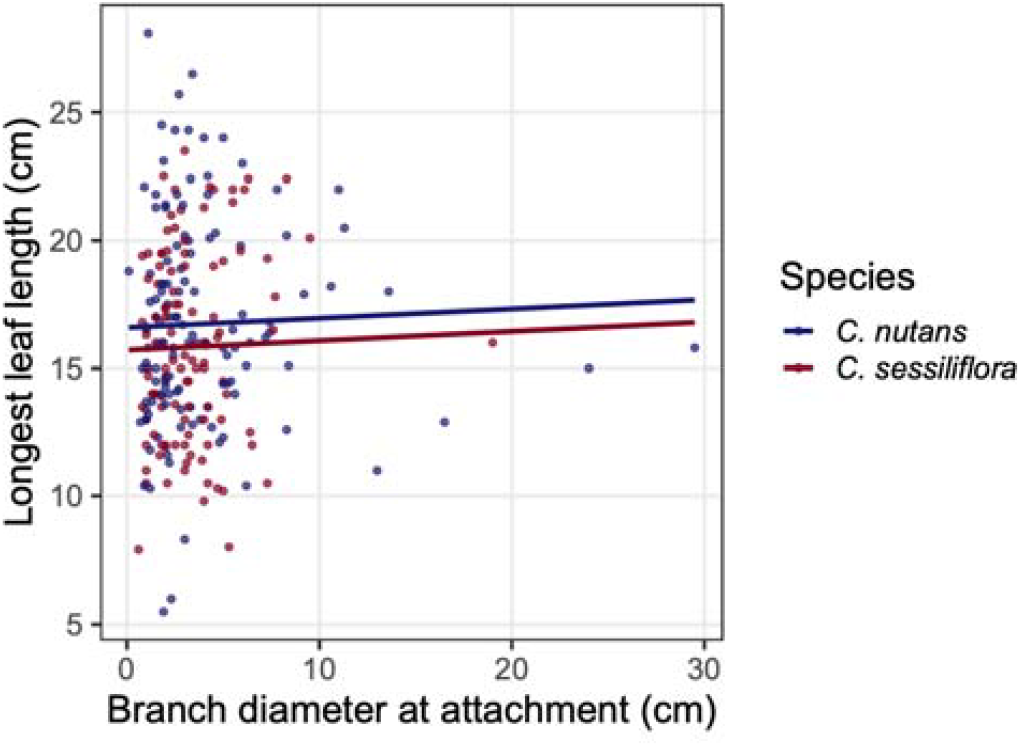
Relationship between host branch diameter and vegetative size in focal *Catopsis* species. Longest leaf length showed little association with branch diameter across sampled individuals of *C. nutans* and *C. sessiliflora*. Lines represent fitted linear models for visualization. These results suggest that substrate diameter alone was not a strong predictor of vegetative size within sampled citrus canopy environments.

**Figure S3.**
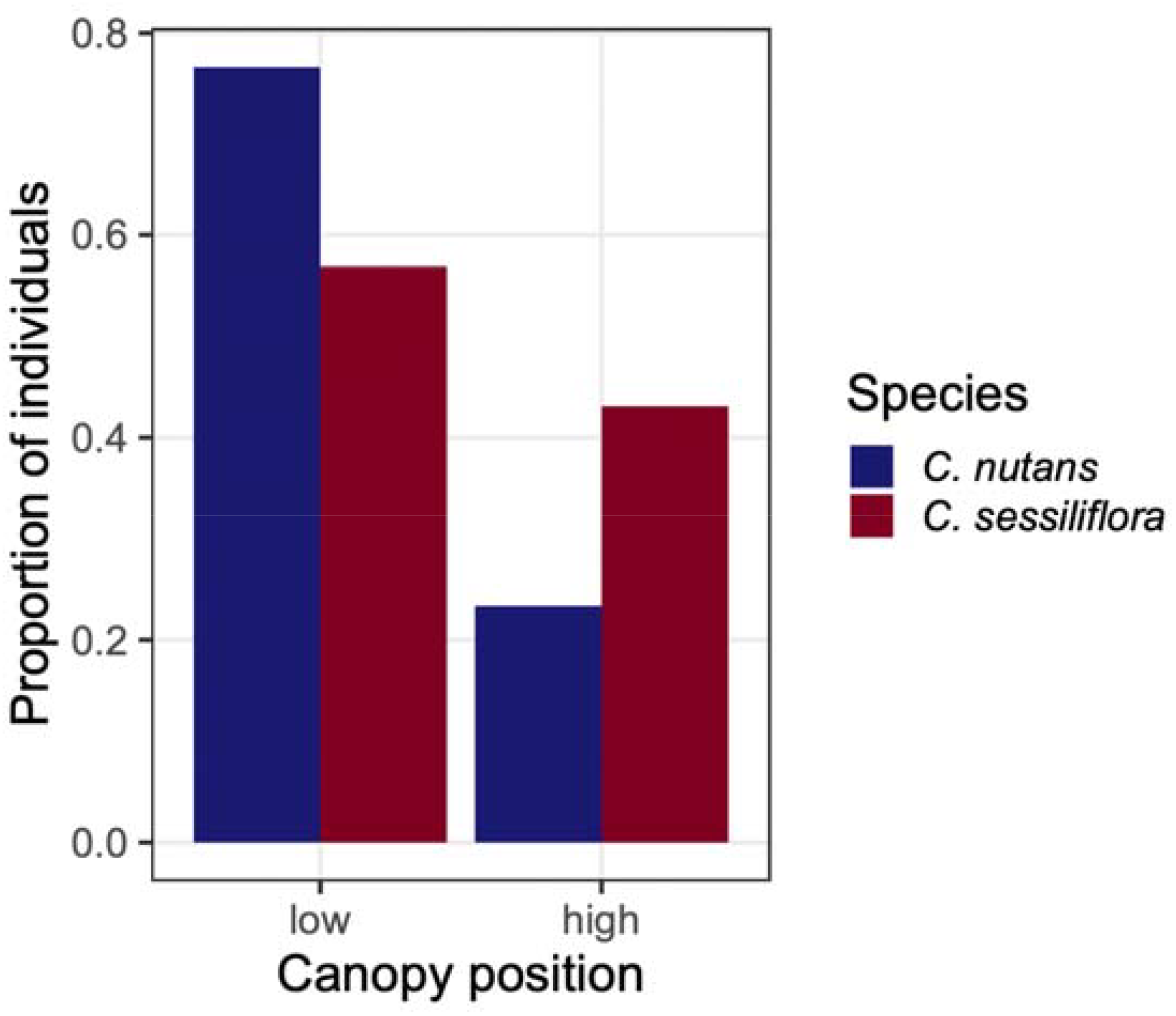
Canopy position occupancy in sympatric populations of *Catopsis nutans* and *C. sessiliflora* occurring within Belize citrus groves. Bars show the proportion of individuals sampled at each canopy position (low, first two primary branches; high, uppermost two branches) within each species. No individuals of either species were recorded at intermediate canopy positions. Both species occurred at both canopy positions, though their distributions differed significantly (Fisher’s exact test: P = 0.017), with *C. sessiliflora* occupying high canopy positions proportionally more often than *C. nutans*.

## Notes

### Competing Interest Statement

The authors have declared no competing interest.

